# EPA-ng: Massively Parallel Evolutionary Placement of Genetic Sequences

**DOI:** 10.1101/291658

**Authors:** Pierre Barbera, Alexey M. Kozlov, Lucas Czech, Benoit Morel, Diego Darriba, Tomáš Flouri, Alexandros Stamatakis

## Abstract

Next Generation Sequencing (NGS) technologies have led to a ubiquity of molecular sequence data. This data avalanche is particularly challenging in metagenetics, which focuses on taxonomic identification of sequences obtained from diverse microbial environments. To achieve this, phylogenetic placement methods determine how these sequences fit into an evolutionary context. Previous implementations of phylogenetic placement algorithms, such as the Evolutionary Placement Algorithm (EPA) included in RAxML, or pplacer, are being increasingly used for this purpose. However, due to the steady progress in NGS technologies, the current implementations face substantial scalability limitations. Here we present EPA-ng, a complete reimplementation of the EPA that is substantially faster, offers a distributed memory parallelization, and integrates concepts from both, RAxML-EPA, and pplacer. EPA-ng can be executed on standard shared memory, as well as on distributed memory systems (e.g., computing clusters). To demonstrate the scalability of EPA-ng we placed 1 billion metagenetic reads from the Tara Oceans Project onto a reference tree with 3,748 taxa in just under 7 hours, using 2,048 cores. Our performance assessment shows that EPA-ng outperforms RAxML-EPA and pplacer by up to a factor of 30 in sequential execution mode, while attaining comparable parallel efficiency on shared memory systems. We further show that the distributed memory parallelization of EPA-ng scales well up to 3,520 cores. EPA-ng is available under the AGPLv3 license: https://github.com/Pbdas/epa-ng

---

In the last decade, advances in genetic sequencing technologies have drastically reduced the price for decoding DNA and dramatically increased the amount of available DNA data. The Tara Oceans Project (Sunagawa et al. 2015), for example, has generated hundreds of billions of environmental sequences. Moreover, sequencing costs are decreasing at a significantly higher rate than computers are becoming faster according to Moore’s law. Therefore, state-of-the art Bioinformatics software is facing a grand scalability challenge.

A common metagenetic data analysis step is to infer the microbiological composition of a given sample. This can be done, for instance, by determining the *best hit* for each query sequence (QS) in a database of reference sequences (RSs), using sequence similarity measures, and by subsequently assigning the taxonomic label of the chosen RS to the QS. However, approaches based on sequence similarity do neither provide, nor use, phylogenetic information about the QS. This can decrease identification accuracy (Koski and Golding 2001), especially when the QSs are only distantly related to the RSs, for example when more closely related QS are simply not available.

*Phylogenetic placement* algorithms alleviate this problem by placing a QS onto a reference tree (RT) inferred on a given set of RSs. This allows for identifying QSs by taking the evolutionary history of the sequences into account. Maximum-Likelihood based phylogenetic placement algorithms have previously been implemented by Matsen et al. (2010) (pplacer) and Berger et al. (2011) (RAxML-EPA). These tools have been successfully employed in a number of studies, for instance, to correlate bacterial composition with disease status (Srinivasan et al. 2012) as well as in diversity studies (Findley et al. 2013; Sunagawa et al. 2013). More recently, we used phylogenetic placements to study protist diversity in rainforest soils (Mahé et al. 2017). In this study we experienced significant throughput and scalability limitations with pplacer and RAxML-EPA. To address them, we re-implemented and parallelized RAxML-EPA from scratch using libpll-2 (Flouri et al. 2017), a state-of-the-art library for phylogenetic likelihood computations.

## Methods

The general algorithm for phylogenetic placement as implemented in EPA-ng, which we call the *placement* procedure, is described in the original paper by Berger et al. (2011). Our supplement also contains a description of the general algorithm.

Like other phylogenetic placement software, EPA-ng operates in two phases: it first quickly determines a set of promising candidate branches for each QS (*preplacement*), and subsequently evaluates the maximum placement likelihood of the QS into this set of candidate branches more thoroughly via numerical optimization routines (*thorough placement*). The user can choose to treat every branch of the tree as a candidate branch, however this induces a significantly higher computational cost. Consequently, by default, EPA-ng dynamically selects a small subset of the available branches via preplacement. Using preplacement heuristics typically reduces the number of thoroughly evaluated branches from thousands (depending on the RT size) to just a handful (depending on the query and reference data).

EPA-ng also offers a second heuristic called *masking* that is similar to the premasking feature in pplacer. It effectively strips the input Multiple Sequence Alignments (MSAs) of all sites that are unlikely to contribute substantially to the placement likelihood score. Such sites consist entirely of gaps, either in the reference or in the query alignment. Additionally, for each individual QS, only the *core* part of the alignment is used to compute the likelihood of a placement. The core of an aligned QS is the sequence with all leading, and trailing gaps discarded. Note that pplacer also discards all gap sites within an individual sequence, including gaps in the core. We opted not to implement this, as our experiments showed that computing these per-site likelihoods, rather than omitting the computations, was more efficient in our implementation.

### Parallelization

EPA-ng offers two levels of parallelism: MPI to split the overall work between the available compute nodes, and OpenMP to parallelize computations within the compute nodes. Such *hybrid* parallelization approaches typically reduce MPI related overheads and yield improved data locality (Rabenseifner et al. 2009).

Figure 1 illustrates how EPA-ng utilizes hybrid parallelism. In hybrid mode, EPA-ng splits the input QS into parts of equal size, such that each MPI-Rank has an equal number of QS to place on the tree. No synchronization is required to achieve this, as each rank computes which part of the data it should process from its rank number and the overall input size.

**Figure 1:**
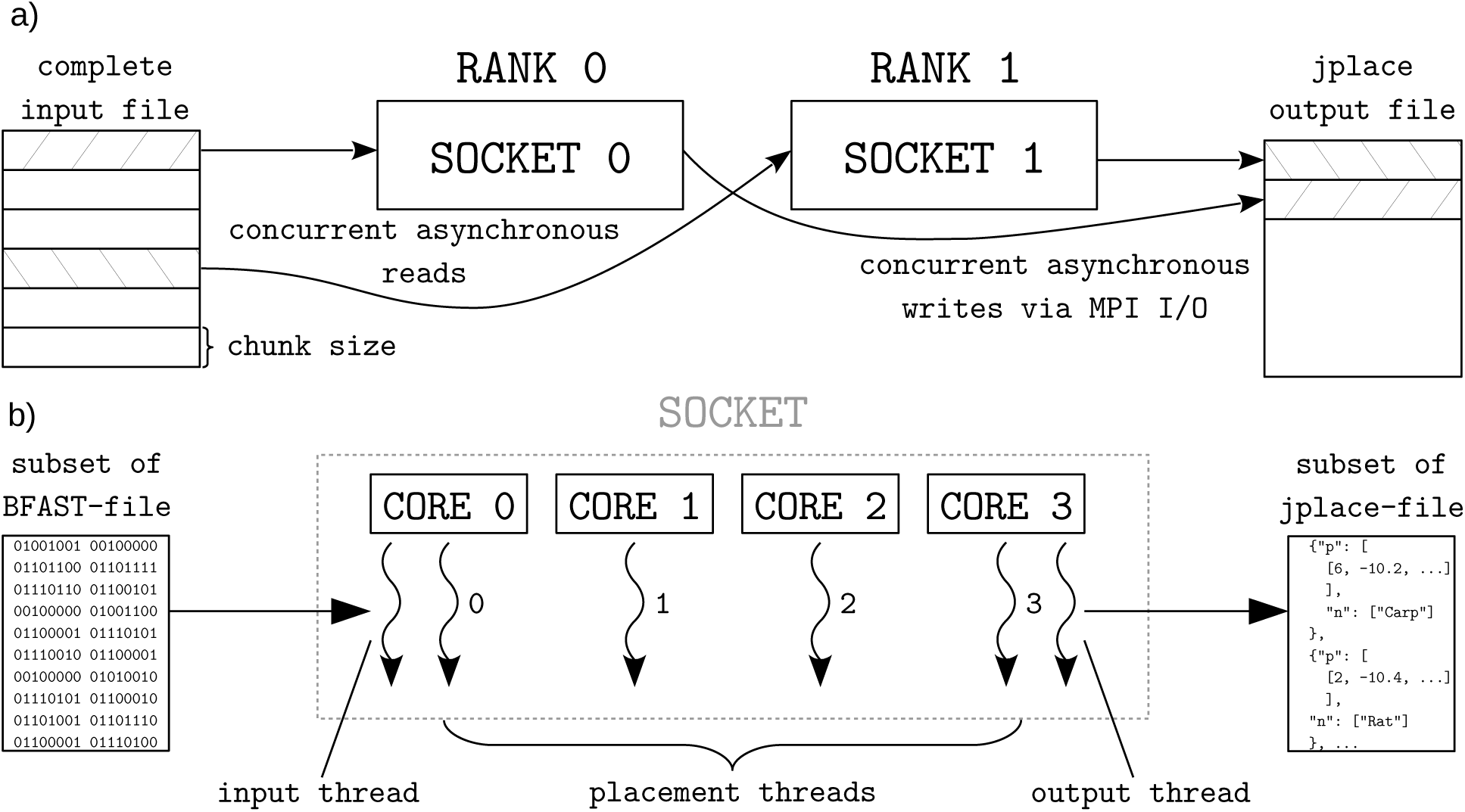
Illustration of the hybrid parallelization scheme implemented in EPA-ng. **a**) shows the parallelization strategy on the level of multiple MPI-Ranks, in this case each assigned to a socket of a node. Each MPI-Rank processes a distinct subset of QS from the input file, and does so in chunks of a given size. When a chunk of QS has been successfully placed the result is written to a global jplace output file, using collective MPI File I/O write operations. **b**) shows the parallelization strategy within each MPI-Rank (in this case: one complete CPU socket). The given subset of the binary input file is read asynchronously by a dedicated input thread, which allows prefetching of one chunk during computation of another. All actual placement work is then split across as many OpenMP worker threads as the user specified (in this case as many as there are physical cores on the socket). Finally, a dedicated output thread writes the per-chunk results to a file, which again allows overlapping of computation and I/O.

For within-node parallelization, we use OpenMP. Here, each thread works on a subset of QS and branches.

## Evaluation

We used three empirical data sets to evaluate and verify EPA-ng, the *neotrop* data set (Mahé et al. 2017), the *bv* data set (Srinivasan et al. 2012), and the *tara* data set (Sunagawa et al. 2015). We compared EPA-ng against pplacer and RAxML-EPA under different settings: with/without masking (not implemented in RAxML-EPA), with/without preplacement. Details on the command line parameters for these distinct settings, as well as full descriptions of the data sets, are provided in the supplement. In the supplement, we also compare the single-node parallel performance parallel efficiency (PE) of the tested programs.

### Verification

In Berger et al. (2011), and Matsen et al. (2010), the authors verify the placement accuracy of their algorithms and implementations via simulation studies and leave-one-out tests on empirical data. As there already exist two highly similar and well-tested evolutionary placement tools, we compare the results of EPA-ng to RAxML-EPAs and pplacers results to verify that our implementation works correctly. We provide a detailed description of the methods deployed for verification and the respective results in the supplement.

Overall, we find that EPA-ng consistently produces results that are situated, on average, closer to the results of RAxML-EPA and pplacer, than the results of RAxML-EPA and pplacer are to each other.

### Sequential Performance

We compared the sequential runtimes of EPA-ng, RAxML-EPA, and pplacer, under two settings. Firstly, with or without the preplacement heuristic. Secondly, with or without the masking heuristic. The combination of these settings results in four distinct comparisons (see Fig. 2). We used 50,000 aligned QSs from the *neotrop* data set, as well as the accompanying reference tree and alignment for this test. The *preplacement* and *masking* settings are as specified in the supplement. Note that RAxML-EPA does not implement masking and therefore respective results are missing.

**Figure 2:**
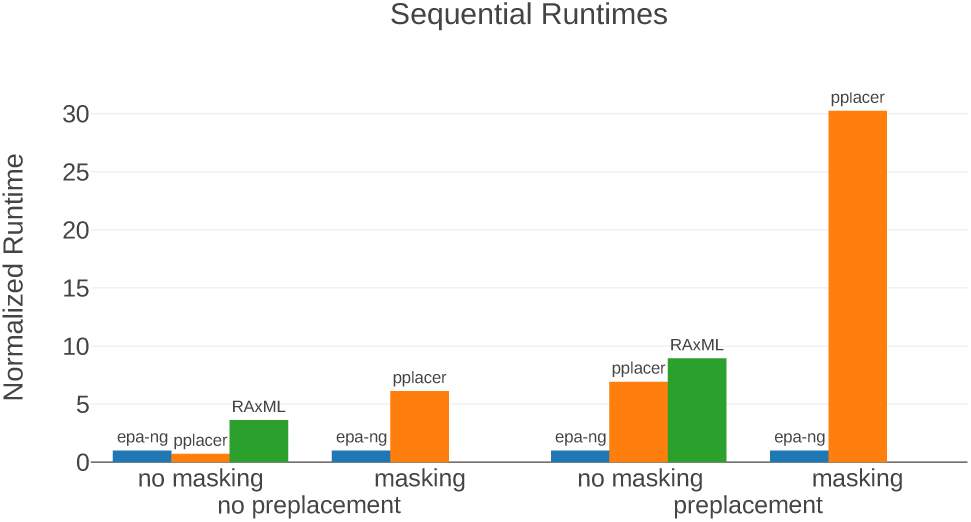
Comparison of sequential runtimes of the three programs EPA-ng, pplacer, and RAxML-EPA, under four different configurations. The y-axis represents the runtime, normalized by the runtime of EPA-ng for each distinct configuration (reads as: *program was x-times slower than* EPA-ng). The absence of data for RAxML-EPA for the masking setting is due to the absence of such a heuristic in RAxML-EPA. The x-axis is scaled logarithmically.

Under most configurations, EPA-ng substantially outperforms the competing programs. The only exception is the case where all heuristics are disabled. In this case we observe a runtime that is ≈ 30% *slower* than for pplacer, while still performing ≈ 3.5-times faster than RAxML-EPA. However, runs with all heuristics disabled do not represent the typical use case. In the configuration using both heuristics, we observe a ≈ 30-fold performance improvement for EPA-ng over pplacer.

### Parallel Performance

We tested the scalability of EPA-ng under three configurations. First, with preplacement and masking heuristics disabled (*thorough* test). Secondly, with only the preplacement heuristic enabled. Lastly, we tested masking in conjunction with preplacement. This corresponds to the default settings (*default* test). Please note that, as RAxML-EPA does not support masking, the respective results are missing.

As runs under these configurations exhibit large absolute runtime differences, we used three distinct input sizes (number of QS) for each of them. The smallest input size for each configuration was selected, such that a respective sequential run terminates within 24 hours. We chose subsequent sizes to be 10, and 100 times, larger, representing medium and large input sizes for each configuration. All scalability tests were based on a set of one million (1M) aligned QSs from the *neotrop* data set. To obtain the desired input sizes, we either sub-sampled (10k, 100k) or replicated (10M, 100M, 1B) the original set of 1M sequences.

As the parallel speedup and the parallel efficiency are calculated based on the fastest sequential execution time, we performed a separate run using the sequential version of EPA-ng (see: *Sequential Performance*). For each configuration, we performed a sequential run for the *small* input volume. As the larger input volumes could not be analyzed sequentially within reasonable time, we multiplied the sequential runtime by 10 and 100, for the medium and large input sizes.

The results are displayed in Figure 3. We observe that the *thorough* test preserves the single-node efficiency (16 cores, ≈ 80% PE) consistently for all core counts and input data sizes. The *preplace* test behaves similarly, but parallel efficiency tends to decrease with increasing core count. This is because the response times are becoming so short, that overheads (e.g., MPI initialization and some pre-computations) start dominating the overall runtime according to Amdahl’s law. This is most pronounced for the 1M QS / 512 cores data point, where PE noticeably declines. The response time in this case was only 83 s, compared to 30,542 s of the corresponding sequential run.

**Figure 3:**
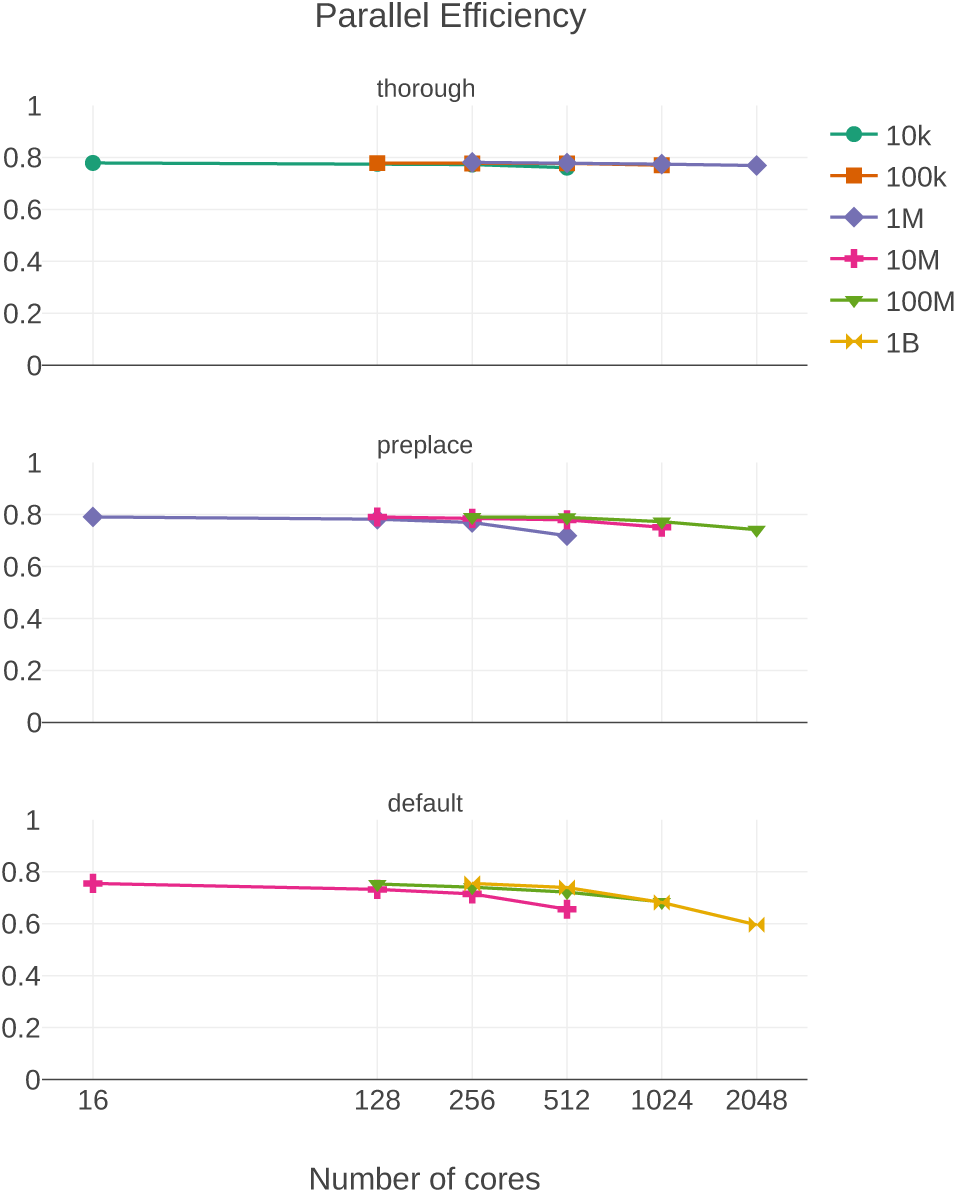
Weak Scaling Parallel Efficiency plot of EPA-ng on a medium-sized cluster. Input files with sizes ranging from ten thousand (10K) to one billion (1B) query sequences. Three different configurations are shown: *thorough*, meaning no preplacement of masking heuristic was employed, *preplace* where only the preplacement heuristic was used, and *default* where both masking and preplacement were employed.

These effects become even more prominent in the *default* run, which shows a PE of ≈ 60% on 2,048 cores. This is primarily due to the increased processing speed when using masking that accelerates preplacement by an additional factor of ≈ 7. As a consequence, operations such as I/O, MPI startup costs, or data pre-processing functions have a more pronounced impact on PE.

## Real-World Showcase

We performed two tests to showcase the improved throughput of EPA-ng and to demonstrate how this enables larger analyses in less time.

### Placing 1 billion metagenomic Tara Ocean sequences

We performed phylogenetic placement of one billion metagenomic fragments (pre-filtered to the 16S rDNA region) against a 3,748 taxa reference tree. Using 2,048 cores (128 compute nodes), we were able to complete this analysis in under 7 hours.

### Extrapolating total reduction in analysis time of Mahé et al. (2017)

We used a representative sample of the *neotrop* data set to obtain runtimes for both, EPA-ng, and RAxML-EPA, using the same settings as in the original study. With this runtime data, we extrapolated the total placement time of the study for both programs. We find that EPA-ng would have required less than half the overall CPU time (RAxML-EPA: 2,173 core days, EPA-ng: 864 core days) under the same heuristic settings (no heuristics). Further, using EPA-ng’s novel heuristics, the placement could have been completed in ≈ 14 core hours (roughly a 3,700-fold runtime reduction).

Our distributed parallelization also improves usability. That is, the user does not have to manually split up the query data (i.e., split the data into smaller chunks which can complete within say 24 hours on a single node) for circumventing common cluster wall time limitations.

## Conclusions and Future Work

In this work, we presented EPA-ng, a highly scalable tool for phylogenetic placement. We showed that it is up to 30 times faster than pplacer and RAxML-EPA when executed sequentially, while yielding qualitatively highly similar results. Moreover, EPA-ng is the first phylogenetic placement implementation that can parallelize over multiple compute nodes of a cluster, enabling analysis of extremely large data sets, while achieving high parallel efficiency and short response times. Our showcase test was executed on 2,048 cores, and placed 1 billion metagenomic query sequences (QSs) from the Tara Oceans project, on a reference tree (RT) with 3,748 taxa, requiring a total runtime of under 7 hours.

We plan to more tightly integrate EPA-ng with upstream and downstream analysis tools, such as programs for aligning the QS against a reference MSA (Berger and Stamatakis 2011), respective placement post-analysis tools (Matsen et al. 2010; Czech and Stamatakis 2017), and methods using the EPA such as SATIVA (Kozlov et al. 2016). In addition, we plan to explore novel approaches for handling increasingly large RTs, such as, for instance, trees comprising all known bacteria.

